# Accounting for small variations in the tracrRNA sequence improves sgRNA activity predictions for CRISPR screening

**DOI:** 10.1101/2022.06.27.497780

**Authors:** Peter C DeWeirdt, Abby V McGee, Fengyi Zheng, Ifunanya Nwolah, Mudra Hegde, John G Doench

## Abstract

CRISPR technology is a powerful tool for studying genome function. To aid in picking sgRNAs that have maximal efficacy against a target of interest from many possible options, several groups have developed models that predict sgRNA on-target activity. Although multiple tracrRNA variants are commonly used for screening, no existing models account for this feature when nominating sgRNAs. Here we develop an on-target model, Rule Set 3, that makes optimal predictions for multiple tracrRNA variants. We validate Rule Set 3 on a new dataset of sgRNAs tiling essential and non-essential genes, demonstrating substantial improvement over prior prediction models. By analyzing the differences in sgRNA activity between tracrRNA variants, we show that Pol III transcription termination is a strong determinant of sgRNA activity. We expect these results to improve the performance of CRISPR screening and inform future research on tracrRNA engineering and sgRNA modeling.

## INTRODUCTION

Pooled screening with CRISPR technology has revolutionized the ease and scale for probing gene function^1,2^. Targeted loci, which are often protein coding genes, can each have hundreds of protospacer adjacent motifs (PAMs), providing many potential single guide RNA (sgRNA) options. It is impractical to assess activity of the millions of potential sgRNAs, so accurate predictions of sgRNA activity are essential for the construction of compact yet potent libraries. Several groups have developed algorithms and web tools to facilitate sgRNA selection^3^.

We previously used a classification model to determine sequence features and developed on-target sgRNA design rules using data from 1,841 sgRNAs (Rule Set 1)^4^. Rule Set 2 then improved upon this initial attempt by incorporating more training data and using a regression model that included additional sequence features^5^. Our guide design portal, CRISPick (broad.io/crispick), has been in continuous operation since 2014 and has averaged 168 design runs per day. Since the development of Rule Set 2, new datasets, features, and model architectures have been developed, which we wanted to include in an updated rule set.

While developing this new rule set, we discovered that small variations in the sequence of the trans-activating CRISPR RNA (tracrRNA) can lead to large differences in activity (here, we will refer to the region of the sgRNA that basepairs to the target DNA as the ‘spacer’ and the remaining structural element of the sgRNA as the ‘tracrRNA’). The majority of published on-target models were trained with sgRNAs that use the tracrRNA as implemented in Hsu et al.^6^, however several other tracrRNAs have been used for screening. Chen et al. modified the Hsu tracrRNA with both a flip – a T to A substitution and compensatory A to T substitution, to disrupt the run of 4 thymidines that can trigger RNA polymerase III termination – and an extension of 5 base pairs in the tetra-loop that is hypothesized to stabilize the sgRNA structure^7,8^. We have also conducted screens with a modification to the Hsu tracrRNA that disrupts the Pol III termination site with a T to G substitution and compensatory A to C substitution, but without any tetra-loop extension (herein named the DeWeirdt tracrRNA)^9^.

To account for the differential effects of these tracrRNA variants, we incorporated tracrRNA identity as a feature in our rule set. Furthermore, to validate our model and gain a deeper understanding for how the different tracrRNAs affect activity, we generated a new dataset with tens of thousands of sgRNAs tiling across essential and non-essential genes for each tracrRNA variant. We show that our updated model, Rule Set 3 (Sequence + Target), makes optimal predictions for all three tracrRNA variants. Additionally, the new screening data demonstrate that disrupting the Pol III termination signal present in the Hsu tracrRNA improves activity for a subset of spacer RNAs, suggesting that the Chen or DeWeirdt tracrRNA may be preferable when target density is a priority, such as base editing screens^10–12^, or when direct detection of the sgRNA is necessary to interpret screening results, such as in some scRNAseq approaches^13^. We expect these results to improve CRISPR-Cas9 screening performance in addition to providing mechanistic insight into tracrRNA-dependent differences in sgRNA activity.

## RESULTS

To understand the current state-of-the-art for on-target modeling, we identified four recently published models^14–17^, and to evaluate these models we collated datasets generated by three different experimental approaches: three genome-wide datasets, four tiling datasets, and four integrated-target datasets^4,5,15,16,18–23^ (**Supplementary Table 1**). Due to the strong interdependence between the models and the collated datasets, we compared the models in a pairwise fashion, allowing us to retain the maximum number of spacer sequences for testing while avoiding leakage between training and testing sets (**Fig 1a**). Spearman correlation was calculated between the observed and predicted activity to assess performance. By this metric, the best performing model was CRISPRon (**Supplementary Fig 1a**).

**Figure 1:**
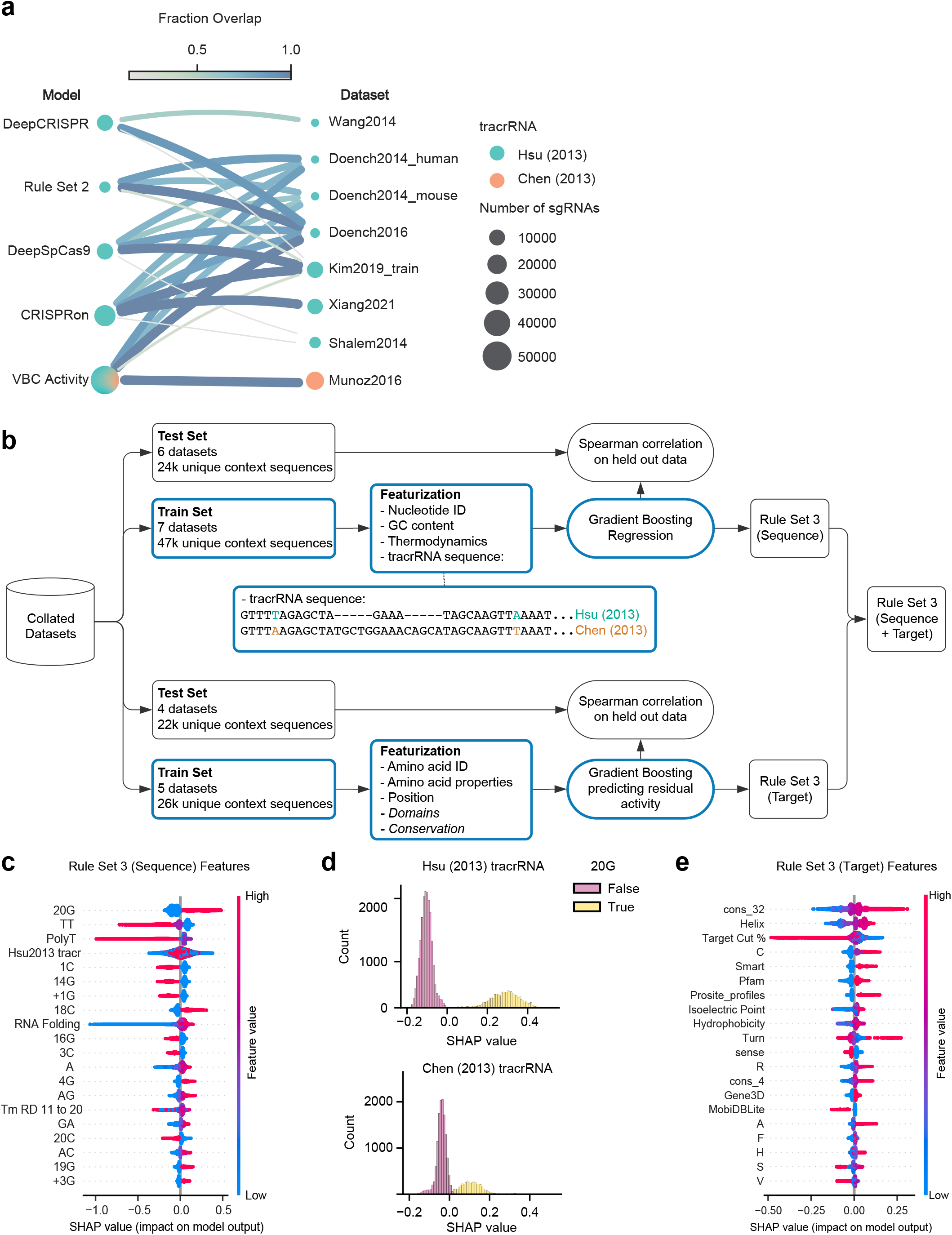
Development of Rule Set 3 (Sequence + Target). a) Fraction overlap between sgRNAs used for training Rule Set 3 and those used for training other on-target models. Edge width and color are proportional to the fraction overlap. Node size is proportional to the number of sgRNAs. b) Schematic depicting Rule Set 3 (Sequence + Target) development. Nucleotide differences in tracrRNA sequences are colored. Models were trained only on the train set, as indicated by blue outlines. Italics indicate features for which information was obtained from existing databases. c) SHAP feature importance for the 20 most important features in Rule Set 3 (Sequence). Each point represents one sgRNA from the training set. Descriptions of model features can be found in Supplementary Table 2. d) Histograms of SHAP values for sgRNAs, colored by guanine status in the 20th sgRNA position and split by tracrRNA identity. e) SHAP feature importance for the 20 most important features in Rule Set 3 (Target). Each point represents one sgRNA from the training set. Descriptions of model features can be found in Supplementary Table 2.

We were surprised that Rule Set 2 marginally outperformed VBC Activity (average difference in Spearman correlation = 0.02), since VBC Activity incorporates Rule Set 2 scores in addition to training on data from Munoz et al. The only dataset where VBC Activity outperformed Rule Set 2 was from Behan et al., which, along with the Munoz data, was one of two collated datasets that utilized the tracrRNA from Chen et al. (**Supplementary Table 1**). This observation led us to hypothesize that there are systematic differences in sgRNA activity that depend on the tracrRNA sequence. To investigate this, we analyzed sgRNA activities from two massive screening efforts, the Dependency Map projects at the Broad and Sanger Institutes^24^. The Broad dataset was screened with the Avana library using the Hsu tracrRNA, while the Sanger dataset was screened with the Human CRISPR Library v.1.0/1.1 and the Chen tracrRNA. These datasets had 876 spacer sequences in common that target essential genes, enabling an assessment of tracrRNA-dependent effects on sgRNA activity. To understand whether there were predictable differences in activity between the tracrRNA variants, we trained gradient boosting models on five splits of the overlapping sgRNAs and measured the Pearson correlation for predictions on held out folds. The average Pearson correlation between the predicted and observed activity differences was 0.34 (**Supplementary Fig 1b**), suggesting that tracrRNA identity should be included as a feature for on-target modeling.

To build a new on-target model that can make optimal predictions for multiple tracrRNA variants, we selected seven datasets for training. This totaled 46,526 unique context sequences, defined as the 20 nucleotide sequence that matches the spacer plus four nucleotides preceding the spacer and six nucleotides (PAM + 3 nucleotides of context) succeeding the spacer RNA; 45% of sequences utilized the Chen tracrRNA. We also held out six datasets for testing (23,629 unique context sequences; 31% with the Chen tracrRNA) (**Supplementary Table 1, Supplementary Data 1**). While convolutional neural networks have proven effective for predicting sgRNA activity^15,16^, we opted for a gradient boosting framework for faster training times^25^. For each sgRNA we encoded the 30mer context sequence using all the features from Rule Set 2 in addition to features to indicate the longest run of each nucleotide in the sgRNA, the melting temperature of the sgRNA:DNA heteroduplex^26^, and the minimum free energy of the folded spacer sequence^27^. We also incorporated categorical variables to indicate which tracrRNA was associated with each spacer, allowing the model to learn features that interact with the tracrRNA (**Fig 1b**). We featurized all of the sgRNAs in the training set and fit a gradient boosting regressor to predict z-scored activity values. We refer to this model as Rule Set 3 (Sequence).

To understand how Rule Set 3 (Sequence) makes its predictions, we calculated Shapley additive explanation (SHAP) values^28^. We found that a G in the tracrRNA-adjacent 20th position of the spacer sequence was the most important feature for activity as has been observed previously (**Fig 1c**)^21^, although there was a strong interaction with the tracrRNA feature, as sgRNAs with the Chen tracrRNA were less affected by the presence of a G in this position (**Fig 1d**). Notably, all three feature classes that were newly added to this model based on prior literature – poly(T), spacer:DNA melting temperature, and minimum free energy – were among the 20 most important features **(Fig 1c, Supplementary Table 2)**. Likewise, tracrRNA identity also proved to be relevant, validating its inclusion in the model. When we considered the held out datasets, Rule Set 3 (Sequence) had the highest Spearman correlation on three of the six datasets, including Behan 2019, which used the Chen tracrRNA (**Supplementary Fig 1c)**. Rule Set 3 (Sequence) predictions were modestly correlated to Rule Set 2 scores for test sgRNAs that used the Hsu tracrRNA (Pearson r = 0.69) (**Supplementary Fig 1d)**.

Several groups have incorporated information about the protein coding sequence, such as protein domains, amino acid sequence, evolutionary conservation, and protein length to predict sgRNA activity^5,17,29^. To test whether these target-site features improve Rule Set 3 (Sequence) scores, we filtered for training data targeting endogenous sites. We calculated the residual activity from Rule Set 3 (Sequence) scores as the outcome variable for the target-site model, Rule Set 3 (Target), ensuring that predictions from the trained model would be additive with Rule Set 3 (Sequence) scores. To test each set of target-site features we split training data into five folds for cross-validation and trained gradient boosting regression models to predict residual activity (**Supplementary Table 1**). Here, target-site features such as amino acid abundance and conservation refer to features of the protein coding region of the gene targeted by the sgRNA and we define activity as the likelihood of disrupting protein function. First, we tested whether protein domains are predictive of sgRNA activity. We queried genes in our training set for functional annotations in Ensembl’s REST API^30^. In total, we obtained functional annotations from 16 sources, including Pfam, Smart, PROSITE, Gene3D, and MobiDB-lite^31–35^. We then tested if the relative abundance of amino acids around the sgRNA cut site was predictive of sgRNA activity. To determine the optimal amino acid window for generating predictions, we tested widths of 2, 4, 8, 16, and 32 amino acids around the cut site. We saw that a width of 16 amino acids led to the highest Spearman correlation with an average value of 0.19 (**Supplementary Fig 2a**). To evaluate the predictive power of evolutionary conservation, we obtained phyloP scores from the UCSC Genome Browser^36,37^. We tested combinations of small and large nucleotide windows around the cut site with the goal of capturing conservation at the cut site as well as more global features such as functional domains. We averaged conservation across 2, 4, or 8 nucleotides for the small window and 16, 32, or 64 nucleotides for the large window. We found that a small width of 4 and a large width of 32 had the best predictive power with an average Spearman correlation of 0.11 (**Supplementary Fig 2b**).

Calculating SHAP values from the trained model, we saw that conservation around the cut site was the most important feature (**Fig 1e**), suggesting that targeting a conserved region of a gene improves sgRNA activity. The second most important feature was the proportion of amino acids around the cut site that are typically found in an alpha-helix (V, I, Y, F, W, L). Michlits et al. noted the favorability of each of these amino acids individually^17^, which we have combined into a single feature. The third most important feature was the relative position of the cut site, where targeting past 85% of the coding sequence led to a steep decrease in sgRNA activity (**Fig 1e**), an effect that has been observed previously^5^. We also examined protein domains^38^; five protein domain sources were among the 20 most important features (Smart, Pfam, PROSITE profiles, Gene3D, and MobiDB-lite), where sgRNAs targeting within an annotated region were more active, with the exception of MobiDB-lite, which identifies long intrinsically disordered regions. Although the relative abundances of seven different amino acids were among the top 20 features, we were unable to identify a biochemical property that explained their importance (**Supplementary Fig 2c**). In fact, the strongest correlate was the fraction of adenine in the codon sequences for an amino acid, potentially indicating that these features were used as a correction to the Rule Set 3 (Sequence) scores after removing a portion of training data that targeted exogenously integrated sites. We tested the target model in combination with the sequence model, which we refer to as Rule Set 3 (Sequence + Target), on held out datasets (**Fig 1b**). Target scores improved the Spearman correlation for all test sets relative to sequence scores alone, with an average improvement of 6.7% (**Supplementary Fig 2d**).

To validate Rule Set 3 and gain mechanistic insight into tracrRNA-dependent differences in spacer activity, we designed a tiling library targeting a subset of essential genes^39^ and non-essential genes^40^. We generated lentiviral vectors that varied in their use of the Hsu, Chen, or DeWeirdt tracrRNA, and each of the three libraries were screened in triplicate in A375 (melanoma) cells stably expressing Cas9. After three weeks, we collected cells, isolated genomic DNA, retrieved the library by PCR, and performed Illumina sequencing to determine the abundance of each sgRNA (**Fig 2a, Supplementary Data 2**).

**Figure 2:**
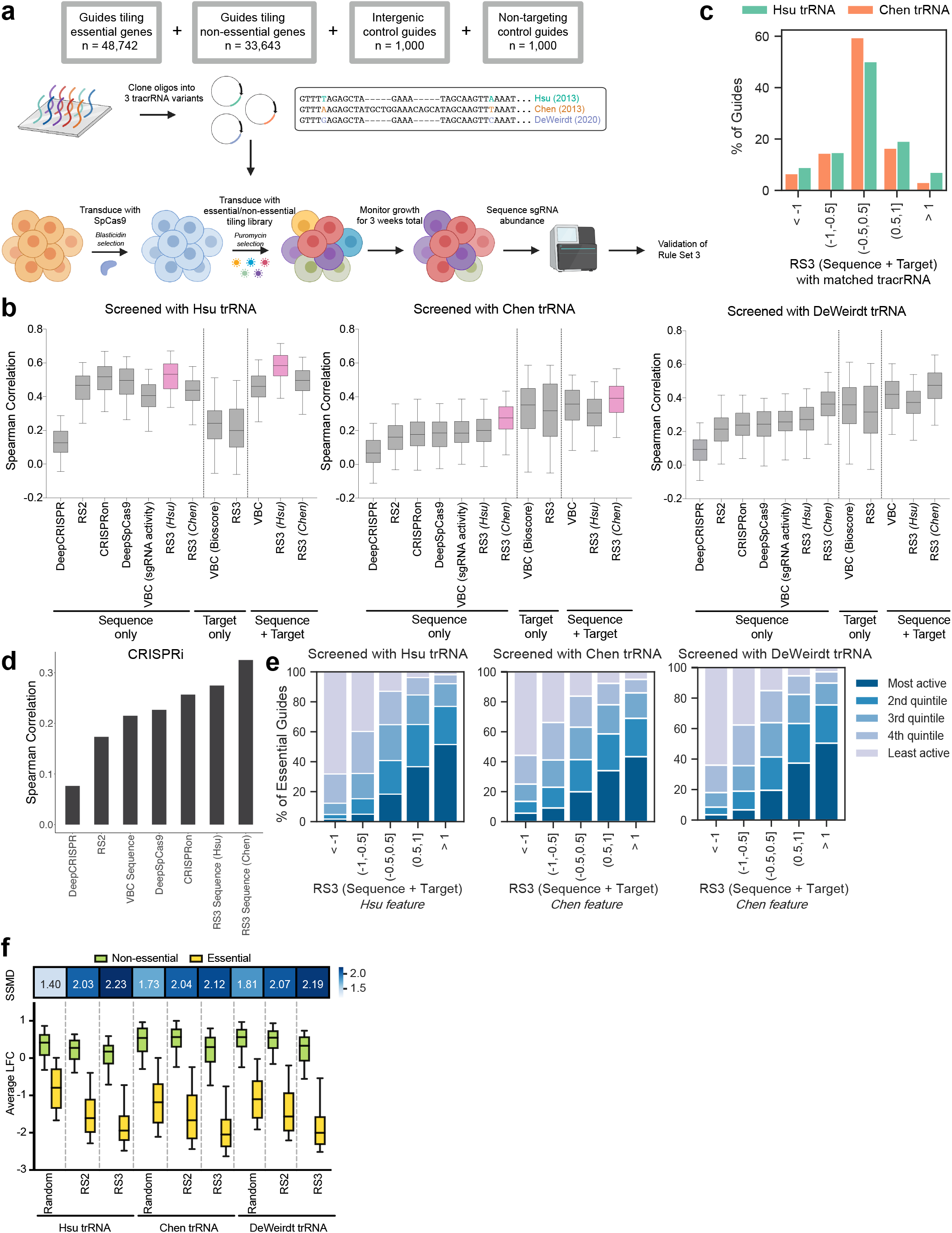
Rule Set 3 (Sequence + Target) validation. a) Schematic depicting essential/non-essential tiling library construction and screening approach. b) Spearman correlations between observed and predicted activity for all essential genes (n=201) in each of the essential/non-essential screens across previous models and Rule Set 3 models. Rule Set 3 models using the same tracrRNA feature as the screen are highlighted in pink. Whiskers show the 5th and 95th percentiles. c) Percent of all sgRNAs targeting essential and non-essential genes broken down by Rule Set 3 (Sequence + Target) bins for the Hsu and Chen tracrRNAs. d) Spearman correlations between predicted scores and the growth phenotype of a tiling CRISPRi dataset across all sequence models. e) Percent of quintiles for sgRNAs from essential genes with at least 20 guides (n=199) binned by Rule Set 3 (Sequence + Target) scores for each screen. Only genes with more than 20 guides are included. f) Average log-fold change of 4 sgRNAs per gene for essential genes (n=201) and non-essential genes (n=198) calculated by picking sgRNAs randomly, using Rule Set 2 or using Rule Set 3 (Sequence + Target) for the tiling library screened with Hsu, Chen, and DeWeirdt tracrRNAs. Rule Set 3 (Sequence + Target) scores used are with the matched tracrRNA. For the screen performed with the DeWeirdt tracrRNA, we used on-target scores with the Chen tracrRNA. Whiskers show 10th and 90th percentile. Heatmap shows the corresponding SSMD scores.

To interpret the results, we first calculated log2-fold-changes (LFCs) compared to the initial library abundance, as determined by sequencing the plasmid DNA (pDNA). As replicates were well correlated (Pearson r = 0.77 - 0.89), we averaged LFCs within each screen (**Supplementary Fig 3a**). Average LFCs across screens were also well correlated, with the Chen and DeWeirdt tracrRNAs showing the highest correlation (Pearson r = 0.89; **Supplementary Fig 3b**). To compare screening performance across tracrRNAs, we calculated the receiver operating characteristic area under the curve (ROC-AUC) defining sgRNAs targeting essential genes as positive controls and non-essential genes as negative controls. The Chen and DeWeirdt tracrRNAs both achieved ROC-AUCs of 0.82, while the Hsu tracrRNA had an ROC-AUC of 0.76 (**Supplementary Fig 3c**). The similarity in performance of the Chen and the DeWeirdt tracrRNAs suggests that the presence or absence of a T in the 5th position has a larger effect on sgRNA activity than the stem extension. We also examined the distribution of various target categories to assess if the controls behaved as expected. In all three screens, the non-targeting controls were tightly distributed and the intergenic controls showed a cutting effect (**Supplementary Fig 3d**).

To evaluate the performance of Rule Set 3 as well as other models on this new dataset, we first removed all spacer sequences that had been seen by any of these models, and then within each essential gene we calculated the Spearman correlation between the predicted scores and the observed sgRNA activity. Rule Set 3 (Sequence + Target) significantly outperformed all other models (t-test p-value < 0.002) when the correct tracrRNA was specified (**Fig 2b, Supplementary Data 3**). We saw better performance when using the Chen tracrRNA as an input to Rule Set 3 when predicting spacers paired with the DeWeirdt tracrRNA, highlighting the importance of disrupting the Pol III termination signal present in the Hsu tracrRNA. Conversely, we saw Rule Set 3 performance decrease when we specified the incorrect tracrRNA. Interestingly, target scores were relatively more helpful than sequence scores for the Chen and DeWeirdt tracrRNAs, although there was large variation across genes, suggesting that sgRNA selection heuristics that over-emphasize target features at the expense of sequence features may lead to poorer performance.

Rule Set 3 (Sequence) had higher Spearman correlations when predicting sgRNAs with the Hsu tracrRNA as opposed to the Chen or DeWeirdt tracrRNAs. This is likely due to a multitude of factors, including a greater diversity of datasets that have the Hsu tracrRNA in the training data, as well as features that are inherently more powerful discriminators for the Hsu tracrRNA. One example of such a feature is having G in the last nucleotide position of the spacer sequence, which has a greater effect on sgRNA activity for the Hsu tracrRNA than the Chen tracrRNA (**Fig 1d**). Further, Rule Set 3 (Sequence) predicts more sgRNAs at the extreme ends of the activity spectrum for the Hsu tracrRNA than the Chen tracrRNA (16% and 9% of sgRNAs with |z-score| > 1 for the Hsu and Chen tracrRNAs respectively; **Fig 2c, Supplementary Data 3**), suggesting the model can identify more discriminating features for the Hsu tracrRNA. We also examined the performance of Rule Set 3 (Sequence) for CRISPRi using a previously published tiling CRISPRi dataset^41^ with sgRNAs targeting the Hart-Moffat essential gene set to assess on-target activity. We calculated the Spearman correlation of the predicted scores and the measured growth phenotype and saw Rule Set 3 (Sequence) performs substantially better than other models, indicating the robustness of Rule Set 3 for predicting other perturbation modalities (**Fig 2d**), although we note that robust CRISPRi predictions should also take into account additional features, such as distance from the transcription start site^42–44^.

To calibrate our understanding of the model outputs, for each essential gene we divided the observed LFCs for each sgRNA into quintiles. We then compared the observed quintiles with predicted activities and saw a direct relationship between the percent of active sgRNAs and Rule Set 3 (Sequence + Target) scores for all tracrRNAs (**Fig 2e, Supplementary Data 3**). For the sgRNAs with the lowest predicted activity, 87.7%, 74.9%, and 82.0%, were observed to be in the lowest or second lowest activity quintile for the Hsu, Chen, and DeWeirdt tracrRNAs respectively. Conversely, for the sgRNAs with the highest predicted activity, 77.0%, 69.0%, and 75.5%, were observed to be in the highest or second highest activity quintile for the Hsu, Chen, and DeWeirdt tracrRNAs, respectively. The large separation between sgRNAs that were observed to be active versus inactive in the highest and lowest predicted bins suggests that sgRNAs picked de novo have a high likelihood of generating gene knockouts for all tracrRNA variants.

To assess how Rule Set 3 (Sequence + Target) impacts library performance in a genome-wide screening context, we simulated a library by picking 4 sgRNAs randomly, using Rule Set 2, or Rule Set 3 (Sequence + Target) with the matched tracrRNA. We observed that across all three screens, as expected, random picking of sgRNAs showed the least separation between average LFCs of essential and non-essential genes (**Fig 2f, Supplementary Data 3**). This separation increased when Rule Set 2 or Rule Set 3 (Sequence + Target) was used to pick sgRNAs. We quantitated this separation by calculating the strictly standardized mean difference (SSMD) between the essential and non-essential genes for each model; a higher SSMD indicates greater separation between the essential and non-essential genes. For all three simulated screens, Rule Set 3 (Sequence + Target) had the highest SSMD. When comparing across tracrRNAs, we found that guides screened with the Hsu tracrRNA and picked using Rule Set 3 (Sequence + Target) had the highest overall SSMD, showing that although there are fewer active sgRNAs with the Hsu tracrRNA (**Supplementary Fig 3c**), by picking a highly active subset of sgRNAs, one can achieve high screening performance.

To understand how each tracrRNA affects spacer activity, we subtracted the z-scores for sgRNAs screened with the Chen tracrRNA from matched sgRNAs screened with the Hsu tracrRNA from the new tiling library. We then trained a gradient boosting model to predict the activity difference from sequence features. We saw that T in spacer positions 17-20 led to relatively lower activity when paired with the Hsu tracrRNA (**Fig 3a, Supplementary Data 4**). On the other hand, G in these same positions led to relatively higher activity for the Hsu tracrRNA. We also observed that spacer RNAs that were predicted to have highly stable secondary structures, as indicated by Gibbs free energy, were less likely to be active when paired with the Hsu tracrRNA.

**Figure 3:**
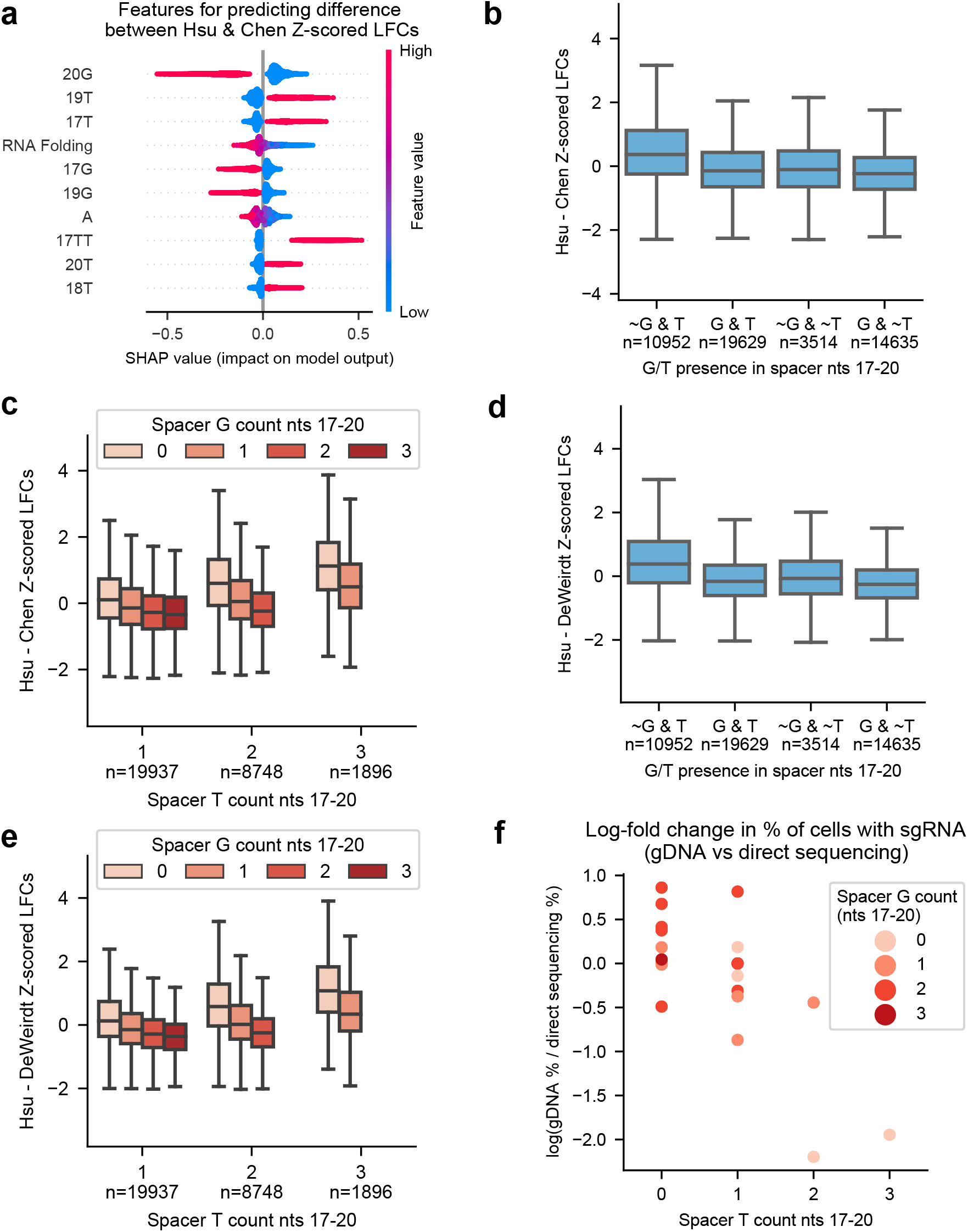
Analysis of differences in tracrRNA activity. a) SHAP feature importance for the 10 most important features for predicting the difference between sgRNAs screened with the Hsu versus Chen tracrRNA. Each point represents one sgRNA from the training set. Descriptions of model features can be found in Supplementary Table 2. b) Box plot of the difference in z-score log-fold changes between sgRNAs screened with the Hsu versus Chen tracrRNA as a function of G/T presence in positions 17-20 of each spacer sequence. The ‘∼’ symbol indicates the nucleotide is not present in this range. c) Box plot of the difference in z-score log-fold changes between sgRNAs screened with the Hsu versus Chen tracrRNA as a function of G/T abundance in positions 17-20 of each spacer sequence. d) Same as (b) but for the Hsu and DeWeirdt tracrRNAs. e) Same as (c) but for the Hsu and DeWeirdt tracrRNAs. f) Data from Mimitou et al. Comparison of sgRNA relative abundance when read out via gDNA or direct sequencing. Spacers are binned by G/T abundance in positions 17-20.

Arimbasseri and colleagues have shown that a stretch of four or more T’s can terminate Pol III transcription, with longer stretches of T’s having stronger effects on transcription termination^45^. The Hsu tracrRNA has a run of four T’s in positions 2-5, whereas the Chen tracrRNA only has three T’s in positions 2-4. In this context, the negative impact that 3’-end T’s have on spacers paired with the Hsu tracrRNA suggests that these spacers are more likely to have a premature termination signal when paired with the Hsu tracrRNA than with the Chen tracrRNA. In support of this hypothesis, a recent study from Graf et al. showed diminished Pol III transcription *in vitro* for two sgRNAs that each had two T’s at the 3’ end of the spacer^46^. Furthermore, they showed that changing the fourth T in the poly-T run of the Hsu tracrRNA to an A restored sgRNA activity.

To investigate whether G’s at the 3’ end of spacer sequences have an attenuating effect on premature transcription termination, we binned sgRNAs by whether they had a G or a T in the last four nucleotides of the spacer. We saw that spacers that had a T and no G had significantly lower activity with the Hsu tracrRNA than spacers that had both a T and G in this region (mean difference = 0.56, p < 0.01; **Fig 3b, Supplementary Data 4**). While spacers with a G and no T were slightly more active than spacers with neither a G nor a T, this difference was smaller (mean difference = 0.16), suggesting that G’s increase the relative activity of spacers paired with the Hsu tracrRNA primarily by inhibiting T-dependent transcription termination signals, as opposed to endowing spacers with some T-independent activity advantage. When we further discretized these bins by the number of T’s and G’s in nucleotides 17-20, we saw that the attenuating behavior of G was sensitive to the number of both T’s and G’s in this region (**Fig 3c, Supplementary Data 4**). We recapitulated these observations when taking the difference in spacer activity between the Hsu and DeWeirdt tracrRNAs (**Fig 3d-e, Supplementary Data 4**), demonstrating that the observed effects are independent of the stem loop extension present in the Chen tracrRNA.

To solidify the connection between 3’ end spacer G/T content and sgRNA expression we analyzed ECCITE-seq data, which captures sgRNA sequence abundance directly^47^. In particular, we analyzed 23 sgRNAs that use the Hsu tracrRNA and were sequenced using both gDNA and direct sgRNA sequencing. We calculated log-fold changes between gDNA and direct sequencing and saw that sgRNA levels significantly decreased as the number of Ts at the end of the spacer increased, controlling for G abundance (linear regression coefficient = -0.53, 95% CI = [-0.848,-0.210], p-value < 0.01; **Fig 3f, Supplementary Data 4**). Conversely, G content tended to enhance transcription, albeit not to a significant level (linear regression coefficient = 0.19; 95% CI = [-0.102, 0.490], p-value = 0.19). Thus, direct sgRNA sequencing supports the hypothesis that use of the Hsu tracrRNA leads to reduced sgRNA expression as a function of G and T prevalence at the end of the spacer sequence.

## DISCUSSION

Here we present Rule Set 3, an optimal model for predicting sgRNA activity for multiple tracrRNA variants. We validate this model on a dataset tiling essential and non-essential genes. By analyzing the differences in activity across multiple tracrRNA variants, we conclude that early Pol III termination is the primary determinant of activity differences between the Hsu and Chen/DeWeirdt tracrRNAs. That a Pol III dependent feature has such a strong impact on sgRNA activity explains a long-standing observation that *in vitro* transcribed sgRNAs used especially in zebrafish systems are poorly predicted by models trained on results from mammalian cell models^48^. Similarly, we expect Rule Set 3 to generalize poorly to sgRNAs being transcribed from a Pol II promoter. As more CRISPR screening data from diverse contexts become available and as transfer learning approaches from machine learning improve, developing models that generalize to a multitude of screening contexts represents a promising direction for future research.

## ACKNOWLEDGEMENTS

We thank all members of the Genetic Perturbation Platform, especially Matthew Greene, Bronte Wen, Doug Alan, Mark Tomko, and Tom Green for software engineering support for CRISPick; the Broad Institute Genomics Platform Walk-up Sequencing group for Illumina sequencing; the Functional Genomics Consortium for funding support; and Priyanka Roy, Audrey Griffith, Zsofia Szegletes, Joshua Dempster, and James McFarland for helpful discussions.

## AUTHOR CONTRIBUTIONS

Conceived of the study: PCD, JGD

Executed genetic screens: AVM, ISN

Performed analyses: PCD, MH, FZ

Created visualizations: PCD, MH, AVM, FZ

Designed libraries: PCD, MH

Curated data: PCD, MH, FZ

Wrote the manuscript: PCD, MH, AVM, FZ, JGD

Supervised the project: JGD

## METHODS

### Vectors

pLX_311-Cas9 (Addgene 96924): SV40 promoter expresses blasticidin resistance; EF1a promoter expresses SpyoCas9.

All guide vectors are derivatives of the lentiGuide vector, with modifications to the tracrRNA. All guide vectors contain the EF1a promoter and puromycin resistance.

pRDA_118 (Addgene 133459): U6 promoter expresses customizable SpCas9 guide with the DeWeirdt (2020) tracrRNA.

pRDA_651: U6 promoter expresses customizable SpCas9 guide with the Hsu (2013) tracrRNA.

pRDA_652: U6 promoter expresses customizable SpCas9 guide with the Chen (2013) tracrRNA.

### Cell lines and culture

A375 cells were obtained from Cancer Cell Line Encyclopedia at the Broad Institute. HEK293Ts were obtained from ATCC (CRL-3216). All cells regularly tested negative for mycoplasma contamination and were maintained in the absence of antibiotics except during screens and lentivirus production, during which media was supplemented with 1% penicillin-streptomycin. Cells were passaged every 2-4 days to maintain exponential growth and were kept in a humidity-controlled 37°C incubator with 5.0% CO2. Media conditions and doses of polybrene, puromycin, and blasticidin were as follows, unless otherwise noted:

A375: RPMI + 10% fetal bovine serum (FBS); 1 μg/mL; 1 μg/mL; 5 μg/mL HEK293T: DMEM + 10% heat-inactivated FBS; N/A; N/A; N/A

### Essential/non-essential tiling library design

201 essential and 198 non-essential genes were randomly chosen from the standard set of essential^40^ and non-essential genes^39^. All possible sgRNA sequences tiling these genes were designed using CRISPick. The library was filtered to exclude any sgRNAs with BsmBI sites or a poly-T sequence. The library was not filtered for promiscuous sgRNAs to enable future efforts focused on off-target analysis using the non-essential genes, but promiscuous guides were excluded from analysis. We also included 1000 controls targeting intergenic sites in the human genome and 1000 non-targeting sgRNAs, resulting in a total library size of 84,609 sgRNAs.

### Library production

Oligonucleotide pools were synthesized by CustomArray. BsmBI recognition sites were appended to each sgRNA sequence (represented here as the run of 20 Ns) along with the appropriate overhang sequences for cloning into the sgRNA expression plasmids. The final oligonucleotide sequence was thus: AGGCACTTGCTCGTACGACGCGTCTCACACCGNNNNNNNNNNNNNNNNNNNNGTTTCGAGACGTTAAGGTGCCGGGCCCACAT.

Primers AGGCACTTGCTCGTACGACG; ATGTGGGCCCGGCACCTTAA were used to amplify the pool using 25 μL 2x NEBnext PCR master mix (New England Biolabs), 2 μL of oligonucleotide pool (∼40 ng), 5 μL of primer mix at a final concentration of 0.5 μM, and 18 μL water. PCR cycling conditions: (1) 98°C for 30 seconds; (2) 53°C for 30 seconds; (3) 72°C for 30 seconds; 24 cycles.

The resulting amplicons were PCR-purified (Qiagen) and cloned into the library vector via Golden Gate cloning with Esp3I (Fisher Scientific) and T7 ligase (Epizyme); the library vector was pre-digested with BsmBI (New England Biolabs). The ligation product was isopropanol precipitated and electroporated into Stbl4 electrocompetent cells (Invitrogen) and grown at 30°C for 16 h on agar with 100 μg/mL carbenicillin. Colonies were scraped and plasmid DNA (pDNA) was prepared (HiSpeed Plasmid Maxi, Qiagen). To confirm library representation and distribution, the pDNA was sequenced.

### Lentivirus production

For pooled library production, 24 h before transfection, 18 × 10^6^ HEK293T cells were seeded in a 175 cm^2^ tissue culture flask in 25 mL of DMEM + 10% heat-inactivated FBS. Transfection was performed using TransIT-LT1 (Mirus) transfection reagent according to the manufacturer’s protocol. Briefly, one solution of Opti-MEM (Corning, 6 mL) and LT1 (305 μL) was combined with a DNA mixture of the packaging plasmid pCMV_VSVG (Addgene 8454, 5 μg), psPAX2 (Addgene 12260, 50 μg), and 40 μg of the transfer vector (e.g. the library pool). The solutions were incubated at room temperature for 20–30 min, then the transfection mixture was added dropwise to the surface of the HEK293T cells. Flasks were transferred to a 37°C incubator for 6–8 h, after which the media was removed and replaced with DMEM + 10% FBS media supplemented with 1% BSA. Virus was harvested 36 h after this media change.

### Derivation of stable cell lines

In order to establish the Cas9 expressing cell line for screens with the essential/non-essential tiling library, A375 cells were transduced with pLX_311-Cas9 and successfully transduced cells were selected with blasticidin for a minimum of 2 weeks. Cells were removed from blasticidin for at least one passage before transduction with the library.

### Pooled screens

For pooled screens, cells were transduced in three biological replicates with the lentiviral library. Transductions were performed at a low multiplicity of infection, using enough cells to achieve a representation of at least 500 transduced cells per sgRNA. We plated cells in polybrene-containing media with 3 × 10^6^ cells per well in a 12-well plate. Plates were centrifuged for 2 h at 931 x g, after which 2 mL of media was added to each well. Plates were then transferred to an incubator for 12-18 h, after which cells were pooled into flasks. Puromycin was added 2 days post-transduction and maintained for 5 days to ensure complete selection for transduced cells. Upon puromycin removal, cells were passaged every 2-3 days for an additional 2 weeks at a minimum of 500x representation, at which point, 21 days post-transduction, cells were collected for subsequent processing. Cell counts were taken at each passage to monitor growth.

### Genomic DNA isolation and sequencing

Genomic DNA (gDNA) was isolated using the KingFisher Flex Purification System with the Mag-Bind® Blood & Tissue DNA HDQ Kit (Omega Bio-Tek). The gDNA concentrations were quantitated by Qubit.

For PCR amplification, gDNA was divided into 100 μL reactions such that each well had at most 10 μg of gDNA. Plasmid DNA (pDNA) was also included at a maximum of 100 pg per well. Per 96-well plate, a master mix consisted of 150 μL DNA Polymerase (Titanium Taq; Takara), 1 mL of 10x buffer, 800 μL of dNTPs (Takara), 50 μL of P5 stagger primer mix (stock at 100 μM concentration), 500 μL of DMSO (if used), and water to bring the final volume to 4 mL. Each well consisted of 50 μL gDNA plus water, 40 μL PCR master mix, and 10 μL of a uniquely barcoded P7 primer (stock at 5 μM concentration). PCR cycling conditions were as follows: (1) 95°C for 1 minute; (2) 94°C for 30 seconds; (3) 52.5°C for 30 seconds; (4) 72°C for 30 seconds; (5) go to (2), x 27; (6) 72°C for 10 minutes. PCR primers were synthesized at Integrated DNA Technologies (IDT). PCR products were purified with Agencourt AMPure XP SPRI beads according to manufacturer’s instructions (Beckman Coulter, A63880), using a 1:1 ratio of beads to PCR product. Samples were sequenced on a HiSeq2500 HighOutput (Illumina) with a 5% spike-in of PhiX.

## QUANTIFICATION AND STATISTICAL ANALYSIS

### On-target modeling

All read count data was transformed to log-fold changes using the poola package (version 0.0.7; https://github.com/gpp-rnd/poola) in Python (version 3.8). For each screen, we selected sgRNAs that were expected to have a phenotype (e.g. sgRNAs targeting essential genes in a viability screen). We filtered any sgRNA that had more than one perfect match in the coding genome. All activities were transformed using the yeo-johnson transformation from scikit-learn (version 0.24.2) and z-scored. Finally, we changed the sign of all activity measurements so the most active sgRNAs had the most positive activity scores. All processed training and testing data can be found on GitHub: https://github.com/gpp-rnd/rs_dev/tree/main/data/processed.

To build Rule Set 3 (Sequence) we used 46,526 unique context sequences from seven datasets. For each sgRNA we encoded the 30mer context sequence using all the features from Rule Set 2 in addition to features to indicate the longest run of each nucleotide in the sgRNA, the melting temperature of the sgRNA:DNA heteroduplex^26^, and the minimum free energy of the folded spacer sequence^27^. We also incorporated categorical variables to indicate which tracrRNA was associated with each spacer, allowing the model to learn features that interact with the tracrRNA. sgRNA features were extracted using the custom Python package sglearn (version 1.2.3; https://github.com/gpp-rnd/sglearn). This package relies on biopython to extract biochemical information about sgRNA sequences^49^.

To fit an optimal gradient boosting model from sequence features, we used the gradient boosting framework from LightGBM (version 3.2.0)^30^ and tuned hyperparameters using Tree Structured Parzen Estimators from Optuna (version 2.7.0)^50^. We tuned the number of leaves (between 8 and 256) and minimum number of samples in a child (between 8 and 256) over 50 hyperparameter iterations. We fixed the learning rate to be 0.01 and used 5,000 boosted trees. All other parameters were kept default. To evaluate each set of hyperparameters we split our dataset into five folds using the StratifiedGroupKFold splitter from scikit-learn. We split the data such that all sgRNAs targeting a gene were either in the train set or the test set for each fold. We also tried to represent each dataset source (e.g. Doench 2016 sgRNAs) proportionally in all of the folds, such that each test set had some sgRNAs from each source. We found that a model with a maximum of 111 leaves per base estimator and a minimum of 199 samples per child performed best. We used these hyperparameters to train our final model.

To build the target model, we used Ensembl’s REST API to query the amino acid sequence around the cut site of each sgRNA (accessed August 9, 2021). We used biopython to get biochemical properties of these amino acid sequences (version 1.79). Ensembl’s REST API was also used to obtain protein domain features. We used the UCSC genome browser’s REST API to get PhyloP conservation scores for each sgRNA (accessed August 9, 2021) ^36,51^. All of these features can be generated in Python using the rs3 package (https://github.com/gpp-rnd/rs3). We tuned hyperparameters for the target model using the same pipeline as Rule Set 3 (Sequence). We found that a model with a maximum of 8 leaves per base estimator and a minimum of 137 samples per child performed best. We used these hyperparameters to train our final model.

To rerun the modeling pipeline, reference the github repository here: https://github.com/gpp-rnd/rs_dev. CRISPick has been updated to incorporate Rule Set 3 (Sequence + Target) scores (broad.io/crispick). Feature importances were calculated using the shap package in Python (version 0.39)^28^.

### Screen analysis

Guide sequences were extracted from sequencing reads by running the PoolQ tool with the search prefix “CACCG” (https://portals.broadinstitute.org/gpp/public/software/poolq). Reads were counted by alignment to a reference file of all possible guide RNAs present in the library. Reads were then assigned to a condition (e.g. a well on the PCR plate) on the basis of the 8 nt index included in the P7 primer. Following deconvolution, the resulting matrix of read counts was first normalized to reads per million within each condition by the following formula: read per guide RNA / total reads per condition x 1e6. Reads per million was then log2-transformed by first adding one to all values, which is necessary in order to take the log of sgRNAs with zero reads.

Prior to further analysis, we filtered out 37 sgRNAs for which the log-normalized reads per million of the pDNA was > 4 standard deviations from the mean in at least one of the screens. We then calculated the log2-fold-change between conditions. All reported LFC values for dropout screens are relative to pDNA. We assessed the correlation between log2-fold-change values of replicates.

We also filtered out sgRNAs targeting essential and non-essential genes that were included in the training set, which constituted ∼6% of all sgRNAs targeting essential genes and ∼0.6% of all sgRNAs targeting non-essential genes.

### SSMD calculation

The strictly standardized mean difference (SSMD) between sgRNAs targetting essential and non-essential genes was calculated using the following formula: 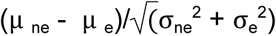 stands for non-essential and *e* stands for essential.

### Data visualization

Figures were created with Python3 and GraphPad Prism (version 9). Schematics were created with BioRender.com.

## DATA AVAILABILITY

The read counts for all screening data and subsequent analyses are provided as Supplementary Data. Fastq files are deposited with the Sequence Read Archive (PRJNA832308).

## STATISTICAL ANALYSIS

All z-scores, Pearson and Spearman correlation coefficients were calculated in Python.

## CODE AVAILABILITY

All custom code used for analysis and example notebooks are available on GitHub: https://github.com/broadinstitute/rs3_manuscript

Code for developing the on-target model can be found on GitHub: https://github.com/gpp-rnd/rs_dev

A python package for scoring sgRNA sequences with Rule Set 3 can be found on GitHub: https://github.com/gpp-rnd/rs3

## FIGURE LEGENDS

**Supplementary Figure 1:**
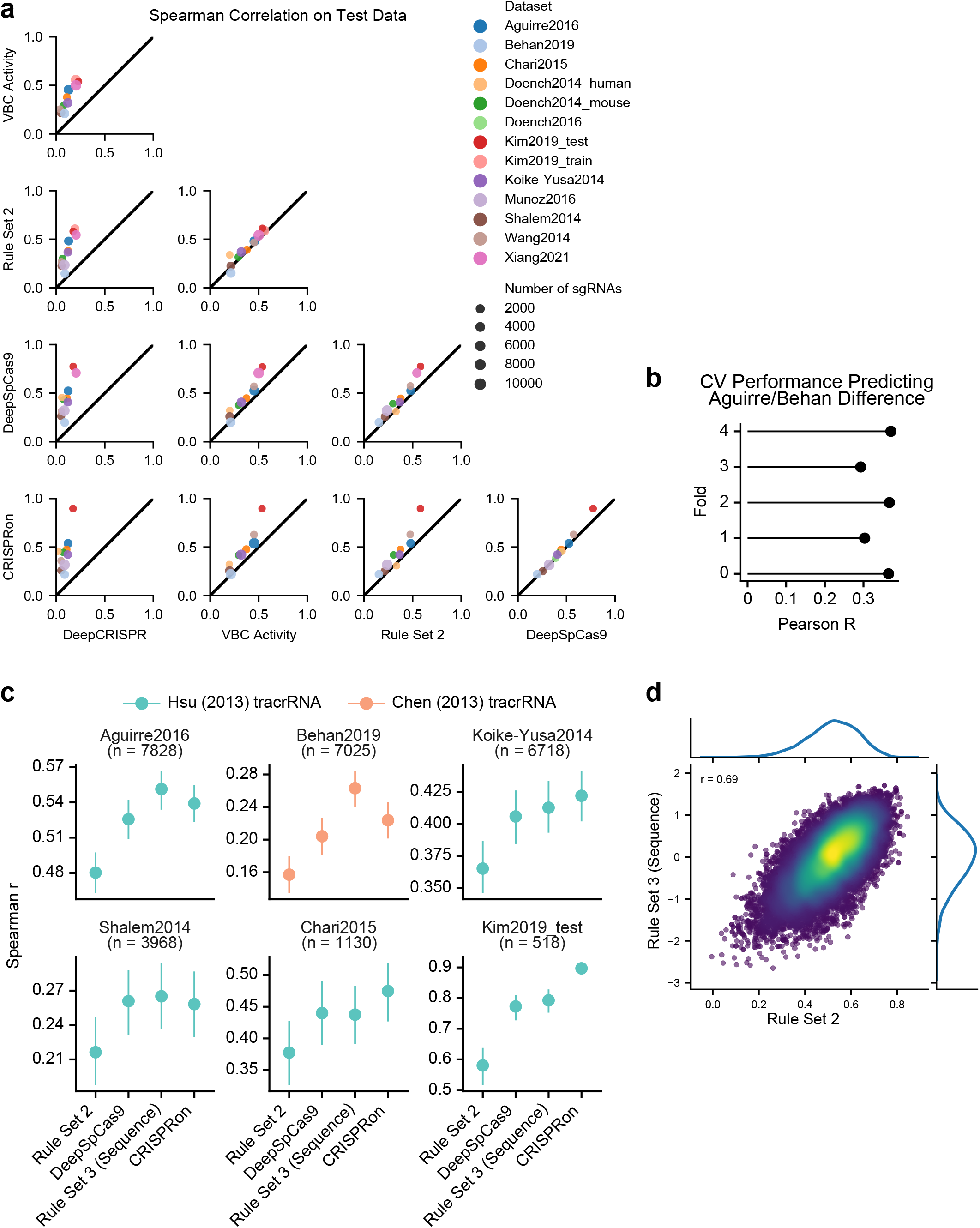
Development of Rule Set 3 (Sequence). a) Spearman correlations between observed and predicted activity for the collated datasets across existing models. b) Pearson correlations on held out folds between the predicted and observed activity differences between sgRNAs in the Behan and Aguirre datasets (CV = cross validation). c) Spearman correlations between observed and predicted activity for six held out datasets across previous models and Rule Set 3 (Sequence). d) Comparison of Rule Set 2 and Rule Set 3 (Sequence) on held out test sgRNAs with Hsu tracrRNA (n=25,268).

**Supplementary Figure 2:**
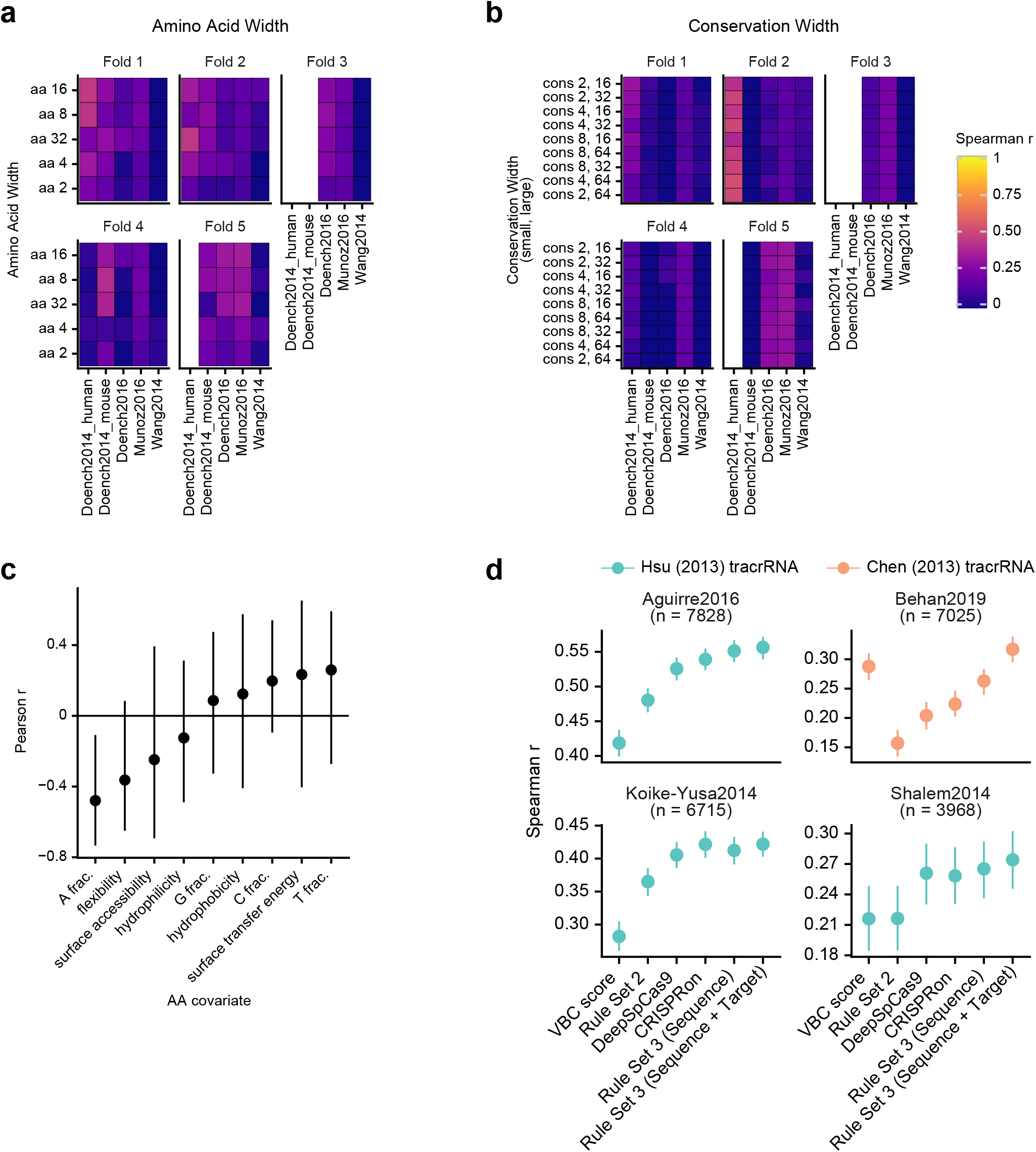
Development of Rule Set 3 (Target). a) Spearman correlations between amino acid window widths around the cut site and cross-validation held out set sgRNA activity. For each fold all sgRNAs targeting a gene were assigned to either the training or testing set and not both. Due to the small number of genes in the Doench 2014 datasets, some folds do not have test sgRNAs from these datasets. b) Spearman correlations between conservation widths around the cut site and cross-validation held out set sgRNA activity. c) Pearson correlations between amino acid biochemical properties and sgRNA activity. d) Spearman correlations between observed and predicted activity for four held out datasets across previous models and Rule Set 3 models.

**Supplementary Figure 3:**
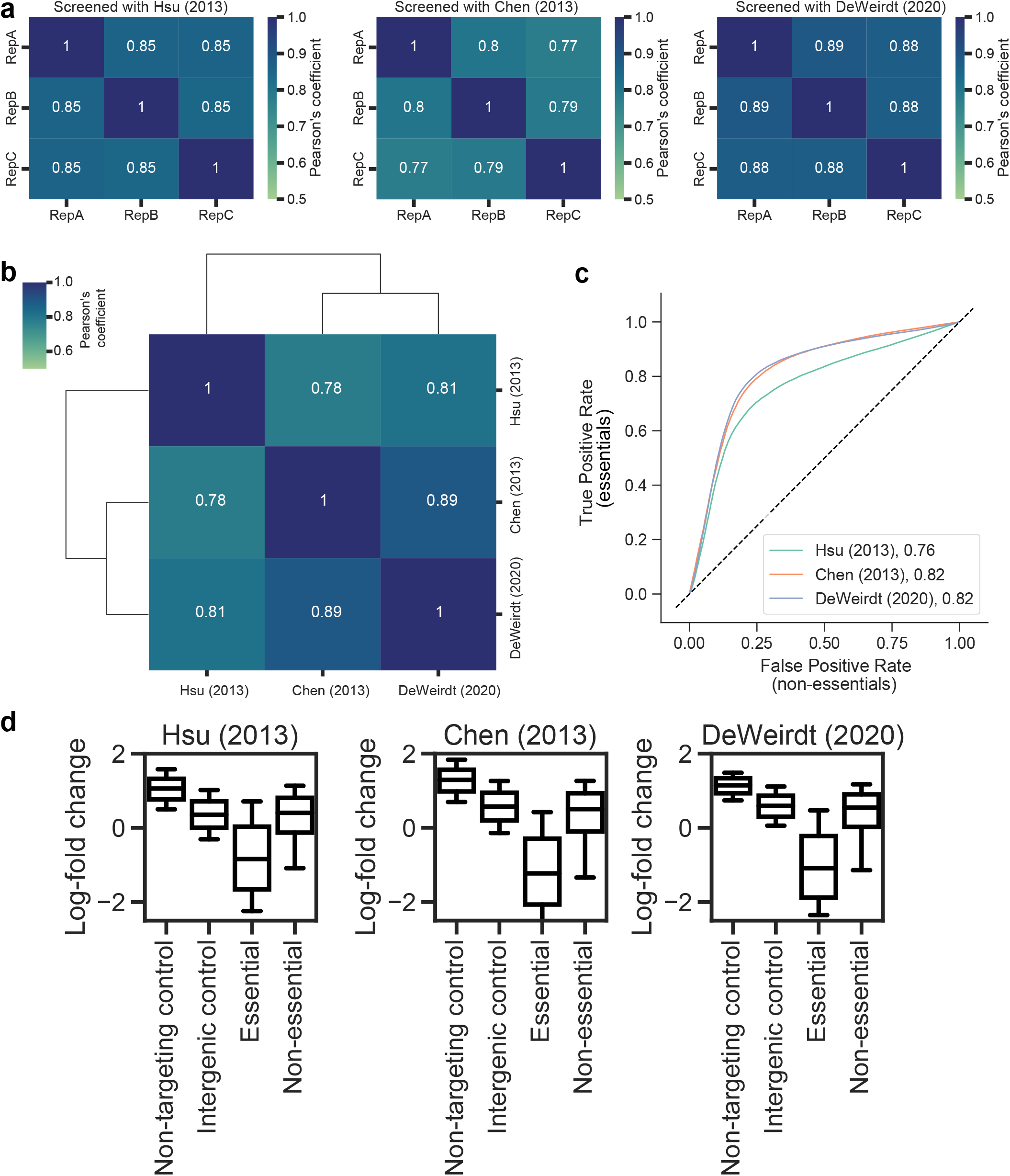
Essential/non-essential tiling library screened with tracrRNA variants. a) Replicate correlation (Pearson’s r) for the essential/non-essential library screened with the three tracrRNA variants in triplicate. b) Correlation (Pearson’s r) between the average log-fold changes for the essential/non-essential library across the three screens. c) ROC plots for the essential/non-essential screen performed with each tracrRNA variant, using sgRNAs targeting essential genes as positive controls and sgRNAs targeting non-essential genes as negative controls. AUC is reported in the graph legend. x=y line is shown. d) Log-fold changes for the different spacer categories in each of the three essential/non-essential screens. Whiskers show 10th and 90th percentile. Number of spacers in each category are as follows: Non-targeting control: 998, Intergenic control: 1000, Essential: 48730, Non-essential: 33621.

## SUPPLEMENTAL DATA

Supplementary Table 1: Compilation of datasets used for training and testing Rule Set 3.

Supplementary Table 2: Description of features used in Rule Set 3.

Supplementary Data 1: Training and testing data for Rule Set 3 (Sequence). **Associated with Fig 1**.

Supplementary Data 2: Essential/non-essential read counts, library annotation. **Associated with Fig 2**.

Supplementary Data 3: On target model Spearman correlations, Rule Set 3 scores for tracrRNA variants, LFCs for Rule Set 2 and Rule Set 3 guide picking, SSMD scores. **Associated with Fig 2**.

Supplementary Data 4: z-score log-fold changes and G/T spacer abundances. **Associated with Fig 3**.

## REFERENCES

1. Doench, J. G. Am I ready for CRISPR? A user’s guide to genetic screens. Nat. Rev. Genet. 19, 67–80 (2018).

2. Przybyla, L. & Gilbert, L. A. A new era in functional genomics screens. Nat. Rev. Genet. (2021) doi:10.1038/s41576-021-00409-w.

3. Hanna, R. E. & Doench, J. G. Design and analysis of CRISPR-Cas experiments. Nat. Biotechnol. (2020) doi:10.1038/s41587-020-0490-7.

4. Doench, J. G. et al. Rational design of highly active sgRNAs for CRISPR-Cas9–mediated gene inactivation. Nat. Biotechnol. 32, 1262–1267 (2014).

5. Doench, J. G. et al. Optimized sgRNA design to maximize activity and minimize off-target effects of CRISPR-Cas9. Nat. Biotechnol. 34, 184–191 (2016).

6. Hsu, P. D. et al. DNA targeting specificity of RNA-guided Cas9 nucleases. Nat. Biotechnol. 31, 827–832 (2013).

7. Chen, B. et al. Dynamic imaging of genomic loci in living human cells by an optimized CRISPR/Cas system. Cell 155, 1479–1491 (2013).

8. Dang, Y. et al. Optimizing sgRNA structure to improve CRISPR-Cas9 knockout efficiency. Genome Biol. 16, 280 (2015).

9. DeWeirdt, P. C. et al. Genetic screens in isogenic mammalian cell lines without single cell cloning. Nat. Commun. 11, 752 (2020).

10. Sangree, A. K. et al. Benchmarking of SpCas9 variants enables deeper base editor screens of BRCA1 and BCL2. Nat. Commun. 13, 1–17 (2022).

11. Hanna, R. E. et al. Massively parallel assessment of human variants with base editor screens. Cell 184, 1064–1080.e20 (2021).

12. Cuella-Martin, R. et al. Functional interrogation of DNA damage response variants with base editing screens. Cell 184, 1081–1097.e19 (2021).

13. Replogle, J. M. et al. Combinatorial single-cell CRISPR screens by direct guide RNA capture and targeted sequencing. Nat. Biotechnol. 38, 954–961 (2020).

14. Chuai, G. et al. DeepCRISPR: optimized CRISPR guide RNA design by deep learning. Genome Biol. 19, 80 (2018).

15. Kim, H. K. et al. SpCas9 activity prediction by DeepSpCas9, a deep learning–based model with high generalization performance. Science Advances 5, eaax9249 (2019).

16. Xiang, X. et al. Enhancing CRISPR-Cas9 gRNA efficiency prediction by data integration and deep learning. Nat. Commun. 12, 1–9 (2021).

17. Michlits, G. et al. Multilayered VBC score predicts sgRNAs that efficiently generate loss-of-function alleles. Nat. Methods 17, 708–716 (2020).

18. Chari, R., Mali, P., Moosburner, M. & Church, G. M. Unraveling CRISPR-Cas9 genome engineering parameters via a library-on-library approach. Nat. Methods 12, 823–826 (2015).

19. Koike-Yusa, H., Li, Y., Tan, E.-P., Del Castillo Velasco-Herrera, M. & Yusa, K. Genome-wide recessive genetic screening in mammalian cells with a lentiviral CRISPR-guide RNA library. Nat. Biotechnol. 32, 267–273 (2013).

20. Shalem, O. et al. Genome-scale CRISPR-Cas9 knockout screening in human cells. Science 343, 84–87 (2014).

21. Wang, T., Wei, J. J., Sabatini, D. M. & Lander, E. S. Genetic Screens in Human Cells Using the CRISPR-Cas9 System. Science 343, 80–84 (2014).

22. Behan, F. M. et al. Prioritization of cancer therapeutic targets using CRISPR–Cas9 screens. Nature 568, 511–516 (2019).

23. Munoz, D. M. et al. CRISPR Screens Provide a Comprehensive Assessment of Cancer Vulnerabilities but Generate False-Positive Hits for Highly Amplified Genomic Regions. Cancer Discov. 6, 900–913 (2016).

24. Dempster, J. M. et al. Agreement between two large pan-cancer CRISPR-Cas9 gene dependency data sets. Nat. Commun. 10, 5817 (2019).

25. Ke, G. et al. LightGBM: A highly efficient gradient boosting decision tree. https://proceedings.neurips.cc/paper/2017/file/6449f44a102fde848669bdd9eb6b76fa-Paper.pdf.

26. Xu, X., Duan, D. & Chen, S.-J. CRISPR-Cas9 cleavage efficiency correlates strongly with target-sgRNA folding stability: from physical mechanism to off-target assessment. Sci. Rep. 7, 143 (2017).

27. Rahman, M. K. & Rahman, M. S. CRISPRpred: A flexible and efficient tool for sgRNAs on-target activity prediction in CRISPR/Cas9 systems. PLoS One 12, e0181943 (2017).

28. Lundberg, S. M. & Lee, S. I. A unified approach to interpreting model predictions. of the 31st international conference on neural … (2017).

29. Schoonenberg, V. A. C. et al. CRISPRO: identification of functional protein coding sequences based on genome editing dense mutagenesis. Genome Biol. 19, 169 (2018).

30. Yates, A. et al. The Ensembl REST API: Ensembl Data for Any Language. Bioinformatics 31, 143–145 (2015).

31. Mistry, J. et al. Pfam: The protein families database in 2021. Nucleic Acids Res. 49, D412–D419 (2021).

32. Letunic, I., Khedkar, S. & Bork, P. SMART: recent updates, new developments and status in 2020. Nucleic Acids Res. 49, D458–D460 (2021).

33. Sigrist, C. J. A. et al. New and continuing developments at PROSITE. Nucleic Acids Res. 41, D344–7 (2013).

34. Lewis, T. E. et al. Gene3D: Extensive prediction of globular domains in proteins. Nucleic Acids Res. 46, D435–D439 (2018).

35. Necci, M., Piovesan, D., Clementel, D., Dosztányi, Z. & Tosatto, S. C. E. MobiDB-lite 3.0: fast consensus annotation of intrinsic disorder flavours in proteins. Bioinformatics (2020) doi:10.1093/bioinformatics/btaa1045.

36. Pollard, K. S., Hubisz, M. J., Rosenbloom, K. R. & Siepel, A. Detection of nonneutral substitution rates on mammalian phylogenies. Genome Res. 20, 110–121 (2010).

37. Karolchik, D. et al. The UCSC Genome Browser Database. Nucleic Acids Res. 31, 51–54 (2003).

38. Shi, J. et al. Discovery of cancer drug targets by CRISPR-Cas9 screening of protein domains. Nat. Biotechnol. 33, 661–667 (2015).

39. Hart, T. et al. High-Resolution CRISPR Screens Reveal Fitness Genes and Genotype-Specific Cancer Liabilities. Cell 163, 1515–1526 (2015).

40. Hart, T., Brown, K. R., Sircoulomb, F., Rottapel, R. & Moffat, J. Measuring error rates in genomic perturbation screens: gold standards for human functional genomics. Mol. Syst. Biol. 10, 733 (2014).

41. Gilbert, L. A. et al. Genome-Scale CRISPR-Mediated Control of Gene Repression and Activation. Cell 159, 647 (2014).

42. Horlbeck, M. A. et al. Compact and highly active next-generation libraries for CRISPR-mediated gene repression and activation. Elife 5, (2016).

43. Radzisheuskaya, A., Shlyueva, D., Müller, I. & Helin, K. Optimizing sgRNA position markedly improves the efficiency of CRISPR/dCas9-mediated transcriptional repression. Nucleic Acids Res. 44, e141 (2016).

44. Sanson, K. R. et al. Optimized libraries for CRISPR-Cas9 genetic screens with multiple modalities. Nat. Commun. 9, 5416 (2018).

45. Arimbasseri, A. G. & Maraia, R. J. Mechanism of Transcription Termination by RNA Polymerase III Utilizes a Non-template Strand Sequence-Specific Signal Element. Molecular Cell vol. 58 1124–1132 (2015).

46. Graf, R., Li, X., Van Trung, C. & Rajewsky, K. sgRNA Sequence Motifs Blocking Efficient CRISPR/Cas9-Mediated Gene Editing. Cell Reports vol. 26 1098–1103.e3 (2019).

47. Mimitou, E. P. et al. Multiplexed detection of proteins, transcriptomes, clonotypes and CRISPR perturbations in single cells. Nat. Methods 16, 409–412 (2019).

48. Haeussler, M. et al. Evaluation of off-target and on-target scoring algorithms and integration into the guide RNA selection tool CRISPOR. Genome Biol. 17, 148 (2016).

49. Cock, P. J. A. et al. Biopython: freely available Python tools for computational molecular biology and bioinformatics. Bioinformatics 25, 1422–1423 (2009).

50. Akiba, T., Sano, S., Yanase, T., Ohta, T. & Koyama, M. Optuna. in Proceedings of the 25th ACM SIGKDD International Conference on Knowledge Discovery & Data Mining (ACM, 2019). doi:10.1145/3292500.3330701.

51. Kent, W. J. The Human Genome Browser at UCSC. Genome Research vol. 12 996–1006 (2002).

